# Contextual associations represented both in neural networks and human behavior

**DOI:** 10.1101/2022.01.13.476195

**Authors:** Elissa M. Aminoff, Shira Baror, Eric W. Roginek, Daniel D. Leeds

## Abstract

Contextual associations facilitate object recognition in human vision. However, the role of context in artificial vision remains elusive as does the characteristics that humans use to define context. We investigated whether contextually related objects (bicycle-helmet) are represented more similarly in convolutional neural networks (CNNs) used for image understanding than unrelated objects (bicycle-fork). Stimuli were of objects against a white background and consisted of a diverse set of contexts (N=73). CNN representations of contextually related objects were more similar to one another than to unrelated objects across all CNN layers. Critically, the similarity found in CNNs correlated with human behavior across three experiments assessing contextual relatedness, emerging significant only in the later layers. The results demonstrate that context is inherently represented in CNNs as a result of object recognition training, and that the representation in the later layers of the network tap into the contextual regularities that predict human behavior.

## INTRODUCTION

Objects do not appear in isolation, but rather embedded within a context. The context of an object includes the regularities of the scene in which it is found, the cluster of other objects it is typically found with, and the spatial relationships between all of these components. These contextual relationships have repeatedly been shown to facilitate human cognition and perception (Biederman et al., 1982; Davenport & Potter, 2004; Koehler & Eckstein, 2015; Lauer et al., 2020; Mudrik et al., 2010; Welbourne et al., 2021). For example, faster reaction times and more accurate responses in recognizing an object are found when the object is either primed by a contextual association (e.g., contextually related scene; Palmer, 1975), or when it is embedded in a congruent context compared with an incongruent context (Biederman et al., 1982; Davenport & Potter, 2004). Thus, contextual associations are a strong cue for understanding our visual world and recognizing objects. However, can visual features alone define context? One way to address what visual regularities are included in context is to examine whether context is represented in artificial visual models.

Building on decades of work, especially in the most recent decade, computer vision has excelled to the level of human performance in recognizing objects (Krizhevsky et al., 2012; Rosenblatt, 1961; Simonyan & Zisserman, 2014). This is largely due to the development of deep convolutional neural networks (CNNs) trained to identify objects in images. CNNs are trained on thousands to millions of images to recognize the statistical regularities that indicate an object’s identity. However, it is unknown whether a CNN inherently learns and utilizes contextual associations to do this, even though it is not explicitly trained to do so. Moreover, if CNNs do learn contextual associations, it is unknown whether these contextual associations relate to the ones utilized in human perception.

Some computer vision work has developed models to integrate scene context into object perception explicitly. Several studies have incorporated context through object spatial position and camera pose, with small to modest improvements in object detection and recognition (Beery et al., 2020; Divvala et al., 2009). Bell et al. (2016) extracted context and object representations through distinct CNNs and merged these two representations to improve object classification nearly twofold, particularly excelling at detecting small objects. These studies suggest that contextual information aids object recognition when context is visually apparent. However, whether context is inherently embedded within the object representation, even when not visually presented, remains an open question. Answering this question is important to understand the similarities and differences between human and computer vision. While multiple studies show that long-term contextual knowledge aids object recognition by human observers, even when objects are presented independent from context, the question remains whether context inherently facilitates computer vision, without being explicitly modeled, and if so, whether context is covertly represented within the representation of objects in a CNN. Extraction of contextual associations during training of artificial object recognition algorithms would further support the functional utility of contextual associations for object recognition. It would also demonstrate that context can be defined through visual regularities and provide a framework for predicting the influence and strength of specific contexts on object recognition.

The current study took a multi-part approach to investigating the role of contextual associations in CNNs and how this role relates to human perception. We focused on examining contextual associations between individual objects (e.g., table-chair) and asked whether these associations are represented in a CNN. To accomplish this, the object representation across the units at each layer of the CNN was first examined. We then looked at whether the representations of contextually related objects (bike - helmet) were more similar to one another than to objects that were unrelated (bike - fork). If so, this would indicate that contextual associations are inherently represented in the network even though the network is not explicitly trained to do so. In order to survey a wide range of potential contextual associations, 73 pairs of contextually related objects were tested (see Figure 1a for examples). Critically, we extracted the CNN representation of objects depicted in images against a white background to prevent confounding the input stimulus with additional contextual information and/or scenery. To address our second question - whether the representations of contextual associations in a CNN were related to the representations that humans use, the contextual similarity represented in a CNN across the related pairs of objects was compared with human performance when rating whether these object pairs were associated with one another. A significant correlation between these measures would indicate the similarity depicted in the CNN was related to contextual associations used in human perception.

**Figure 1:**
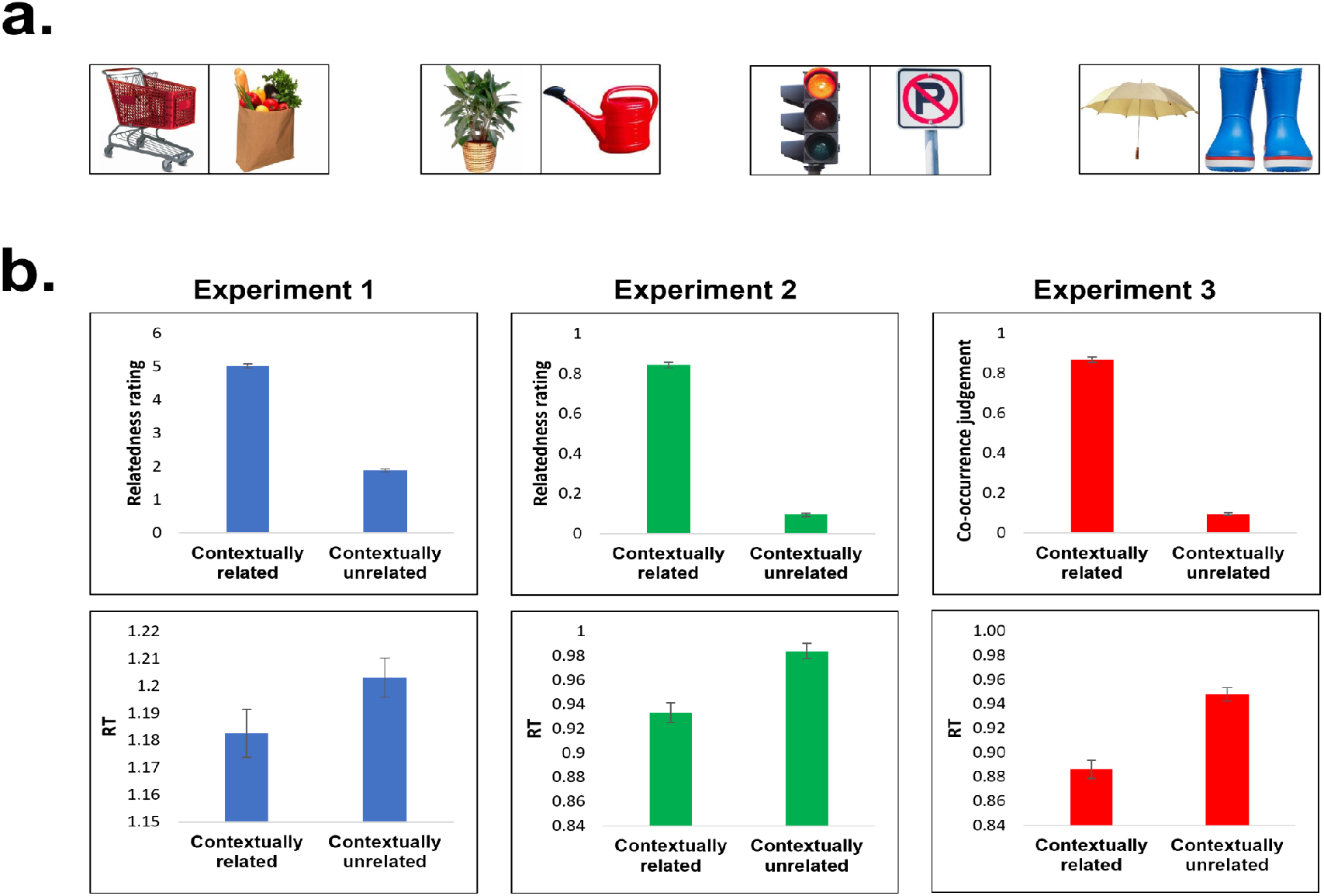
**a)** Examples of the stimuli used. 73 contexts were chosen, each with 2 paired objects. All pictures were photographs of objects against a white background. **b)** Behavioral validation results. Across three experiments, see Methods, contextually related objects are judged as related significantly more than unrelated objects (top row). This is reflected in reaction time as well, such that related judgements of contextually related objects are made significantly faster than judgements of contextual unrelated objects. Error bars represent standard error of the mean.

## RESULTS

### Validation of stimuli

Contextual associations between objects were first assessed in three human behavioral experiments. In these experiments, two images belonging to two object categories (e.g., an easel and a palette) were presented simultaneously and participants were asked to judge whether the objects were related. In all experiments, 73 pairs of contextually related object categories were assessed. To make sure judgements generalize beyond exemplar-specific attributes, for each object category (146 object categories, comprising the 73 pairs) five different exemplars of the object were employed. For example, five different pictures of hairdryers and five different pictures of barber chairs comprised the two object categories that as a pair form the barber context. In the experiments, each context was represented across four different trials: two trials in which both objects were contextually related; and two trials in which the objects were unrelated. Unrelated pairs consisted of swapping object categories from the contextually related pairs (e.g., hairdryer - bird). Relatedness was assessed using three different tasks while presenting two objects simultaneously, side by side: In Experiment 1 (N = 32), relatedness was assessed on a 6 point scale, such that each pair of objects was rated from unrelated (1) to related (6). In Experiment 2 (N = 20), a two-alternative-forced-choice was employed, asking participants to press the button (s) if the objects were related, and another button (d) if the objects were unrelated. In Experiment 3 (N=20), participants pressed a button (s) if they expected to find the two objects in the same picture, and another button (f) if they did not. For all three tasks, participants were asked to respond as quickly and as accurately as they could.

Results validated that contextually related objects were indeed more strongly related to one another than unrelated objects (see Figure 1b). In Experiment 1, contextually related object categories were rated as more related (mean 5.02) than unrelated object categories (mean 1.88; t(72) = 44.48, p < 4.09×10^−54^). Participants were also faster at rating the contextually related objects as related (mean 1.18 seconds) compared with the unrelated object categories (mean 1.2 seconds; t(72) = -2.09, p < 0.04). These results replicated across the two additional experiments. In Experiment 2, when tasked with a two alternative forced choice, participants rated contextually related object categories faster and as more related (mean response: 84% related responses, reaction time (RT): 0.93 seconds) compared with contextually unrelated pairs of object categories (mean response: 9% related responses, RT: 0.98 seconds; paired response t-test: t(72) = 47.22, p < 6.35×10^−56^; paired RT t-test: t(72) = -6.28, p < 2.20×10^−8^). In Experiment 3, when asking whether the objects belonged in the same picture, related object categories were more predicted to appear at the same picture (mean response: 86% same picture) and were responded to faster (mean RT = 0.88 seconds) compared with unrelated objects (mean response: 9% same picture, mean RT: 0.94 seconds; paired response t-test: t(72) = 55.59, p < 6.91×10^−61^; paired RT t-test: t(72) = -7.6, p < 8.29×10^−11^). Thus, given our behavioral data we can confirm that humans do perceive the contextually related objects as strongly related to one another compared with the unrelated objects.

### Neural network representation of context

Would a CNN network also treat contextually related pairs of objects as related, even though it was not explicitly trained to do so? To test CNN context-integration, the representation of an object across units in a layer was compared to the representation of a contextually related object and to the representation of an unrelated object. Stronger similarity to the contextually related object compared to the unrelated object would indicate that contextual associations were included in the object’s representation. We first tested context-integration with a popular benchmark CNN, VGG 16 (Simonyan & Zisserman, 2014), trained on image recognition with the ImageNet dataset (Deng et al., 2009). To provide a framework in which to interpret the results of the contextually related pairs, we first compared the representations of objects that belong to the same category. This comparison demonstrates similarity of representations in the CNN based on object category membership (see Figure 2), as the network was explicitly trained to link together objects in the same category. As mentioned above, each object was depicted in five different exemplars. To look at categorical representation, the similarity of the CNN representations across the different exemplars of the same category was evaluated. Similarity is measured as the ratio of pairwise similarities of stimuli within a category (bird 1 vs bird 2) versus the similarities outside of the category (bird 1 vs palette 1). Ratios above 1 would indicate that there is greater similarity across categorically related objects than unrelated objects; thus, we expected similarity ratios above 1. Results showed that objects that are categorically related to one another (e.g., two hairdryers) are represented more similarly in every layer of the CNN studied above the first layer (Figure 2c). Ratios were consistently above 1.1 starting at layer 4 (t(141) > 14.8, p < 2×10^−29^) and grew more significant with each subsequent layer (the output layer of the network was not included in the analysis). The maximum similarity was found in the last layers (ratios 2.23 and 3.05, in layers 25 and 27); this heightened similarity was expected since a more invariant representation of category is thought to be represented higher in the network.

**Figure 2:**
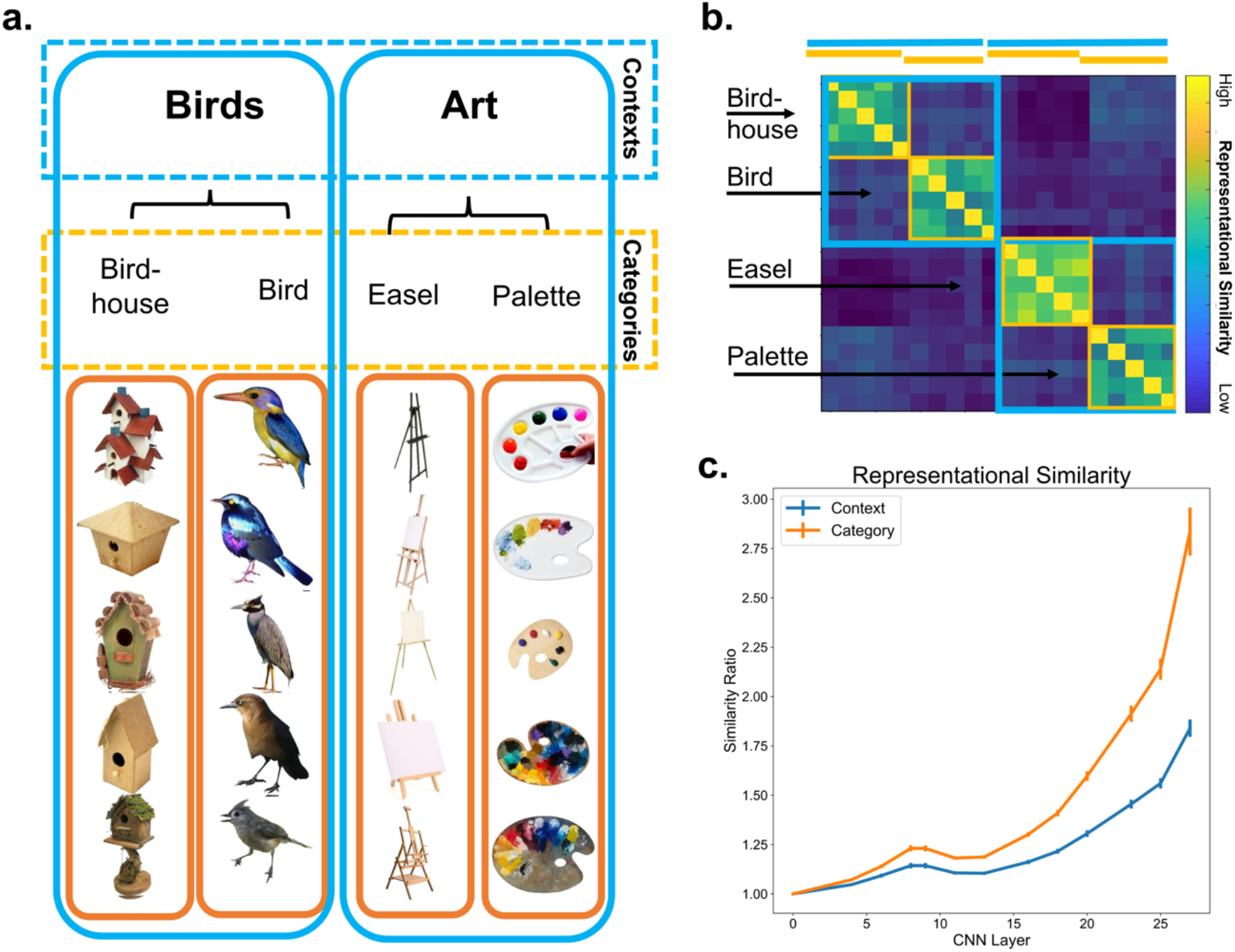
**a)** Examples of the contrasts of interest. Five different exemplars of each object were used. Category similarity was assessed by comparing representations in the CNN across exemplars of the same category (orange outlines). Context similarity was assessed by comparing the representations of exemplars across the object paired categories (e.g., one easel and one pallet exemplar, blue outlines). **b)** Example of similarity matrix for each comparison. **c)** Representational similarity of images in the same category (orange) and same context (blue) at each layer of VGG16 CNN. Similarity computed as ratio of average in-group versus out-group similarities for each group. Error bars represent standard error of the mean.

Next, in comparison to categorical relationships, contextual relationships were investigated. These relationships are not explicitly trained but are rather implicitly learned based on statistical regularities found in the training stimuli. We found contextually related pairs of objects were represented more similarly in the network (despite not looking like one another, e.g., lamp-chair) than unrelated pairs (e.g., lamp-stroller). Two contextually related pairs had maximum contextual similarity ratios several magnitudes above the mean. These were considered outliers and were removed from subsequent analyses to prevent false inflation of the results. Thus, all analyses henceforth included 71 contexts. Surprisingly, the level of context-based similarity was significant at every layer studied above the first layer. Ratios were consistently above 1.01 and significantly above 1 starting at layer 4 (t(70) > 8.5, p < 3×10^−12^). The magnitude and significance of the ratio grew with each subsequent layer, reaching the maximum similarity in the last hidden layers of the network (ratios 1.62 and 2.01, in layers 25 and 27). Naturally, the degree of context similarity was less than the similarity exhibited for categorical relationships (Figure 2c), as would be expected given the network is trained to provide explicit output identifying the object category. Nevertheless, although the network was trained to recognize individual object categories, the network also implicitly represented contextual associations between objects.

### Correlation between human behavior and neural network representation

The critical test in our analysis was to determine whether the representation of contextual associations in a CNN had any relation to human behavior. To address this, Pearson correlations between the behavioral performance and the similarity between the CNN representations of contextually related objects were computed. Specifically, behavioral performance, both relatedness judgements and reaction time, were correlated with the maximum similarity ratio value representing the contextual relatedness between pairs of objects in a CNN. The maximum ratio was found at the highest hidden layer (layer 27) in all but 1 of the 71 context pairs (the defiant context’s maximum ratio was found at the penultimate layer, layer 25). This correlation was computed separately for each behavioral experiment (see Figure 3). The purpose of looking at all three experiments was to reflect the ability to replicate the findings across tasks and independent sets of participants. The resulting analysis found a correlation between the level of relatedness as indicated by human performance and similarity in the CNN network. This was significant in the second experiment (r(69) = .23, p < .05), and marginally significant in the second and third experiment (Ex. 1: r(69) = .21, p < .08; Ex. 3: r(69) = .20, p < .1). The positive direction of the correlation demonstrated the more participants found the pair of objects related to one another, the more similar the objects were represented across units in a CNN (Figure 3a). Furthermore, to use a more stringent test, the more implicit and sensitive measure of reaction time was compared with the similarity of representation in a CNN. The resulting correlation demonstrated a significant negative correlation in experiments 1 and 2 (Ex. 1: r(69) = -.24, p < .05; Ex. 2: r(69) = -.27, p < .02). The negative correlation demonstrated that the faster participants were able to judge the relatedness of the objects, the more similar contextually related objects were represented in the CNN (Figure 3b). This supports the proposal that contextual associations are represented in a CNN, and those representations are related to human behavior.

**Figure 3:**
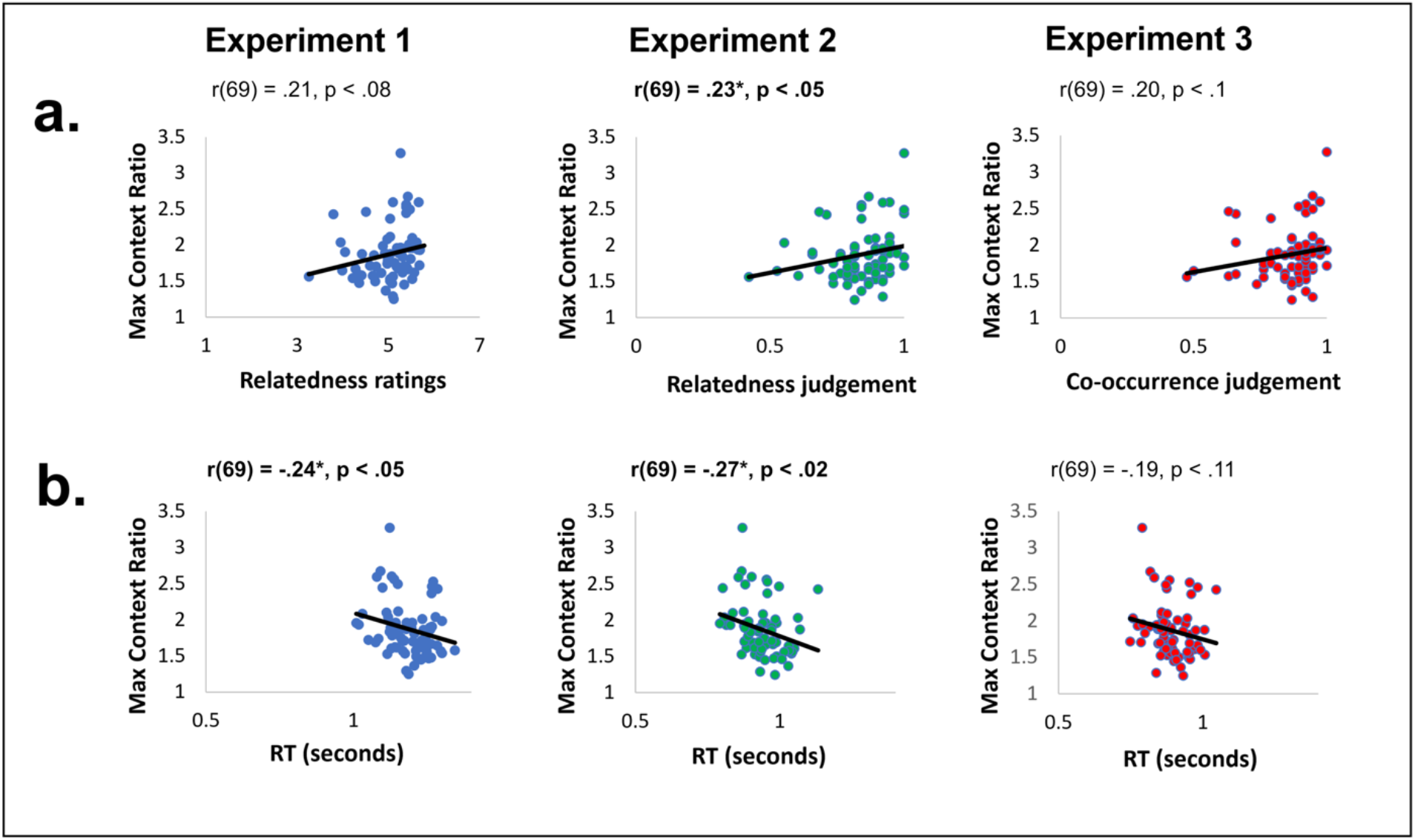
correlations between the representation of context in VGG16 and contextual performance in humans. **a**) Positive correlations between human ratings of contextual relatedness and context similarity in the CNN; significant in Experiment 2, marginal correlation in Experiment 1 & 3. **b)** Negative correlations between human reaction time and context similarity in the CNN; significant in Experiment 1 & 2. Correlations are computed over 71 contexts.

The hierarchical nature of the layers (lowest layer 0, and highest hidden layer 27) in a CNN can also provide additional insight into understanding when the similarity of contextually related objects in a CNN is most relevant to human behavior. To find out which layers were most related to human behavior, we correlated behavioral responses of the contextually related trials with the similarity of the contextually related object representations in each extracted VGG 16 layer (N = 13). Correlation across the different layers was only computed for those comparisons that demonstrated a significant correlation with the maximum similarity ratio discussed above (Experiment 1 RT, Experiment 2 response and reaction time). Correlations were significant only in the later layers of the network (see Table 1). Significance between reaction time and similarity of CNN representations first appeared in layer 20 (Experiment 1 RT: r(69) = -.25, p < .03, Experiment 2 RT: r(69) = -.27, p < .03) and continued to be significant through layer 27 (Experiment 1 RT: r(69) = -.24, p < .05, Experiment 2 RT: r(69) = -.27, p < .02). Significance between relatedness responses and similarity of CNN representations was only significant at the last hidden layer, layer 27 (Experiment 2 relatedness response: r(69) = .23, p < .05).

**Table 1:**
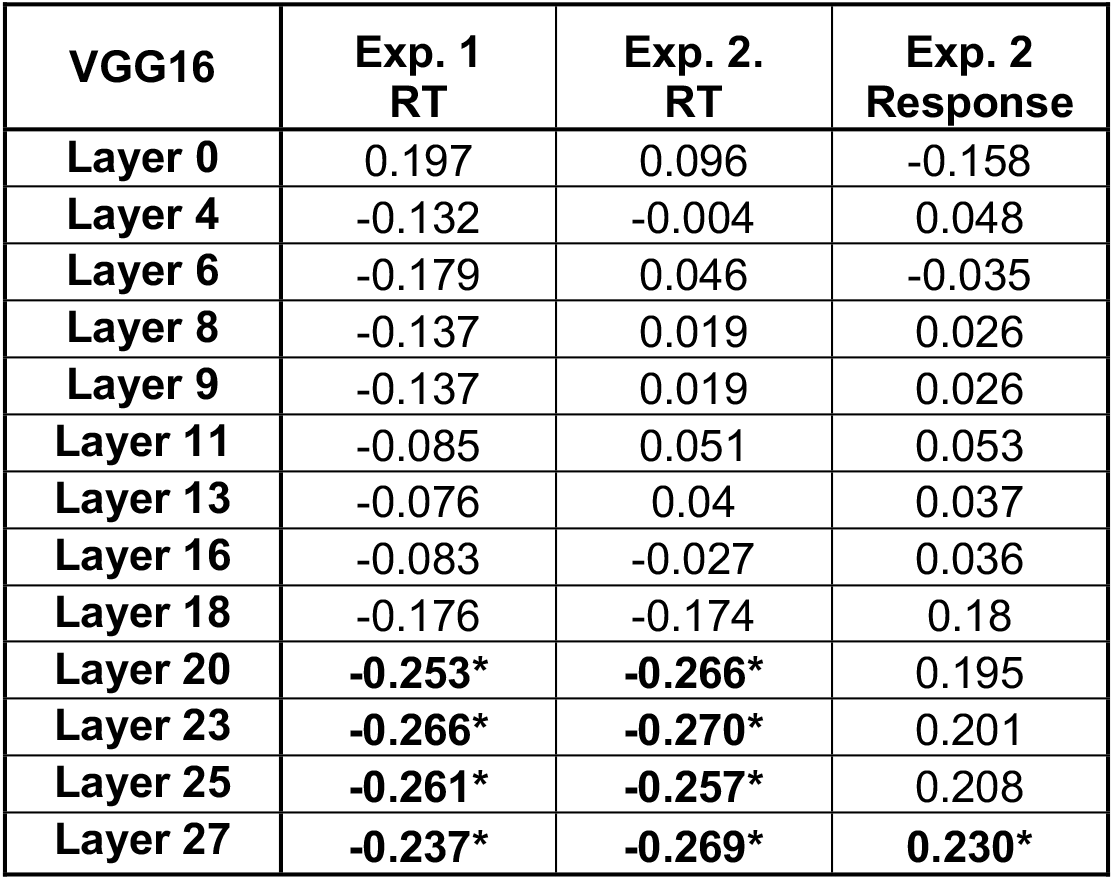
Pearson correlation between human behavioral performance and similarity in the VGG16 CNN network layers. Pearson r values between behavior (Experiment 1 reaction time, Experiment 2 reaction time, and Experiment 2 relatedness response) are listed in each cell. Cells marked with * and bold indicates correlations that were significant at p < .05. Significance of the correlation was only found in the later layers of VGG16.

This analysis revealed that although contextually related objects have a significantly similar representation across almost all layers of the CNN, only those representations in the later layers are related to the contextual associations that humans use.

### Exploratory analysis of context representation in other CNNs

Now that we established that contextual associations are represented in one CNN, we investigated whether these associations were unique to the VGG16 network, or whether contextual associations were represented in a variety of CNNs. We therefore studied CNNs that varied by number of layers, by whether they had a recurrent structure, and by whether they were trained on ImageNet (i.e., object based) or Places365 (i.e., scene based). Similar to the procedure carried out for VGG16, for each network, in-group versus out-group similarity ratios were measured across 71 Contexts and 142 Categories at the maximum ratio layer, comparing these ratios to the null hypothesis (ratio = 1, Figure 4). The results show that the representation of contextual associations exists significantly in each network studied (p’s < 2×10^−23^). For full results, please see Supplemental Materials Figure 1. Some trends were found, however, differentiating between networks. Non-recurrent networks showed substantially higher in-out ratios for both category and context than did recurrent networks (t(70) = 21.4, p < 3×10^−32^), suggesting simpler representations are more effectively fit to object and context properties. ImageNet-trained networks showed higher in-out context ratios than did Places365-trained networks (t(70) = 23.6, p < 6×10^−35^), suggesting that the representational similarity of objects from the same context is more effectively captured when learning to distinguish objects rather than when learning to distinguish overall scenes.

**Figure 4:**
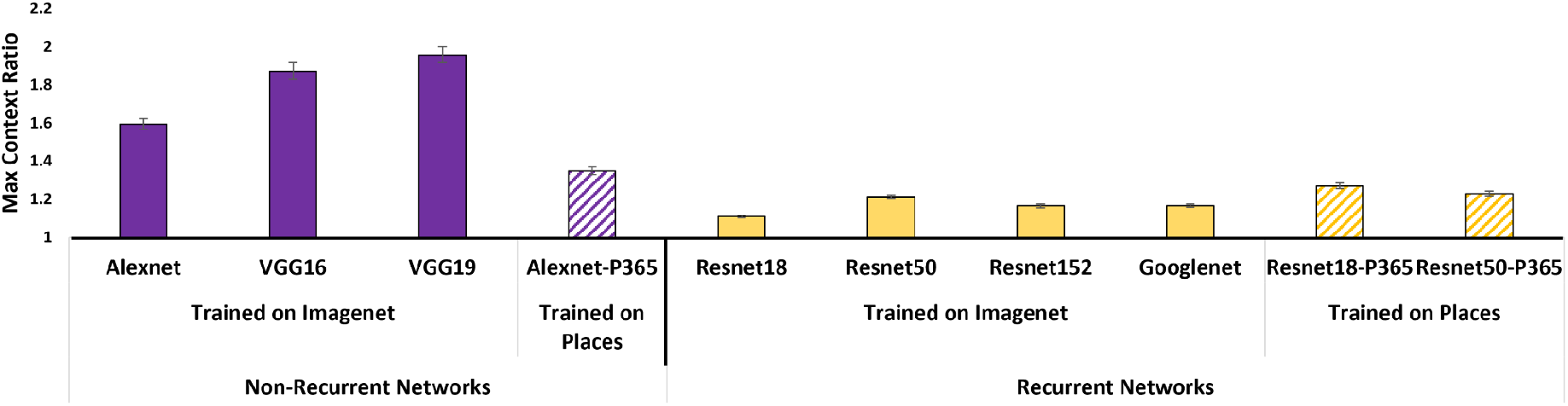
Contextual representations across a variety of networks. Highest similarity ratio for ten CNN architectures trained on ImageNet (solid bars) and Place365 (striped bars) data sets. Each network tested demonstrated a significant effect of context (p’s < 9×10^−24^). This effect is significantly greater in non-recurrent networks (purple bars) compared with recurrent networks (yellow bars).

## DISCUSSION

This study used a CNN framework to examine whether contextual associations between objects are inherently extracted while learning to perform object classification, and whether the CNN-learned representational space for objects and contexts was related to human behavior. Our findings revealed that objects that were contextually related to one another were more similarly represented at each layer of the CNN compared to unrelated counter objects. This indicates that commonalities related to context are integrated in the object CNN representation. Thus, despite being dissimilar in their visual features (e.g., umbrella and rainboots) and despite being presented on a white background with no visual context depicted, the network recognized a commonality between contextually related pairs of objects (bike helmet - bike) compared with unrelated objects (e.g., bike - football). This finding confirms that contextual associations are inherently coded in a CNN that is designed to recognize objects, despite it not being explicitly trained to capture context. More importantly, we found that the extent to which the representations of contextually related objects were similar in the CNN correlated with human performance, both with how fast human participants responded in judging whether these object pairs were related, as well as with the degree of relatedness judgment itself.

Our study focused on the VGG16 network, a non-recurrent network trained on ImageNet with the purpose of recognizing objects within an image, ultimately identifying each image as one of a possible 1000 objects. The network’s successful optimization for object categorization is evident in the high similarity in the representation of objects that share the same category (e.g., two different birdhouses) compared with objects that don’t share the same category. The significant similarity between objects of the same category was evident across all studied layers of the network and, as expected, increased with each subsequent layer. This analysis of category-based similarity was used as control and validated the use of this parameter as evaluating the representations encapsulated in the network. We then used it as a benchmark to compare the representation of context in the network, and found that belonging to a shared context, not only to a shared category, increased object representational similarity in the network as well.

Furthermore, in our investigation of context it was critical to present the objects against a white background in order to prevent confounding the presented stimulus with any visual contextual information (distinguishing from other recent related work, e.g., Bracci et al., 2021). Even though objects were presented in isolation, the similarity related to a shared context was evident in every layer of the network after the first layer, increasing with each subsequent layer. As later layers are more tuned to the ultimate goal of object categorization (Rafegas et al., 2020), it is very possible that achieving the goal of object categorization relies on contextual information with every additional layer. As expected, due to the targeted task of the network, the similarity based on object category was significantly higher than the similarity based on context. However, our results suggest regularities associated with context were correlated with object categories, and thus were integrated into the object representations to facilitate category recognition. This relation was true not only for VGG 16, but for each of the nine other CNNs that were tested. Thus, both non-recurrent and recurrent networks inherently integrated context within object representations, as well as networks that were trained on both ImageNet and Places365. Interestingly, those networks trained to recognize scene categories (i.e., those trained on Places365) did not have significantly stronger context representations, suggesting that context is important for scene categories, but also critical to the task of object recognition itself. Previous studies, as highlighted in the introduction, were able to improve the network’s performance by explicitly modeling context. Our results are novel in showing that even without explicit training, context is inherently captured in the network. Context, therefore, is not just evident in facilitating human perception, but also facilitates artificial perception in computer vision.

The degree of similarity in the CNN representations of contextually related objects correlated with human behavior both with regards to rating the degree to which two objects are related, as well as with how quickly these ratings were made. Hence, the degree of similarity in a CNN can be used to predict context effects on human behavior, such as the strength and weight a particular context can have on facilitating object recognition. More generally, this also supports that the context represented in a CNN is relevant to understanding and modeling how context is derived in human behavior. Context facilitation of human perception is typically framed as utilizing prior knowledge to generate predictions and expectations on our perception in a top-down manner. In a CNN, however, the computations that are applied to determine the representation at each layer of the network are driven by bottom-up statistical regularities found in the trained images. The results of the current study support the idea that contextual relations between objects are derived bottom-up from the statistical regularities of co-occurrence, which is supported by previous neuroimaging results (Aminoff & Tarr, 2015). Thus, the correlation to human behavior supports a model that signifies at least some of the contextual associations that humans utilize in perception derive from the statistical regularities one experiences in the environment.

However, statistical co-occurrence alone is not the only requirement. For example, we found that the computer-human correlation was more strongly evident in Experiments 1 and 2, where participants were asked to make general relatedness judgments, compared with Experiment 3 where participants were specifically instructed to rate how likely the objects were to be found in the same picture. If the correlation between human and computer vision reflected simple statistical co-occurrences between items in an image, we would expect the highest correlation to emerge in Experiment 3 in which we targeted co-occurrence of objects. In addition, interestingly, the significant correlations between CNN and human behavior were only found in the later layers of the network, even though context was significantly represented at almost each layer of the CNN. Thus, relying just on statistical regularities to explain context in human cognition is too simplistic. For example, visual regularities represented at lower levels may represent features such as lines and edges – but these did not correlate with human performance. Only those statistical regularities emerging at higher levels of visual processing in the network, which may represent more complex and invariant representations, were relevant to human performance. Further research is needed to uncover the differences of context representation at low levels versus high levels of a CNN. This will provide more insight into how context is derived, utilized, and updated in humans.

The results of this study demonstrate the importance of taking into account contextual associations in models of object recognition and image understanding in both human and artificial vision. And indeed, recent work incorporates context more readily in the deep networks used for image understanding. For example, scene graphs have played an important role in recent computer vision approaches to modeling static and dynamic scenes; co-occurring objects in a given image are recognized and characterized through their relations with one-another (Ost et al., 2021; Xu et al., 2017; Yang et al., 2018; Zhang et al., 2020). However, patterns of inter-object association are not explicitly tied to the underlying scene in each image, and scenes can vary with inter-object association. These models that incorporate a cluster of contextual information are a promising avenue for further work in biological and computational studies as they demonstrated the strong role for context in modeling vision.

Computer vision can achieve human level performance in recognizing objects of a scene (e.g., He et al., 2015), however, there is still a gap between how humans understand an image and how computer vision understands an image. This difference may be fueled by a divergence in learning experiences – humans assemble visual and contextual knowledge across a lifetime of linearly evolving experiences that build off one another, while CNNs typically train on a static training set of images mixed together in less determined order and more limited in overall diversity. It is possible that further context learning may be key to bridging the gap between the biological and artificial systems, and that the more neural networks utilize context in ways similar to those of humans, for example, relying more on the context representation at later layers, the more computer vision may understand unusual or unique images in the same way humans effortlessly do. Ultimately, the more we bridge the gaps between human and artificial vision, the more computer vision can be applied to aiding and working in concert with human vision, for example supplementing and aiding people with visual impairments.

In conclusion, our study shows that context is inherently encapsulated in neural networks, and that this correlates with human perception. Thus, object recognition trained CNNs represent context, emphasizing those contexts with the most impact on human cognition. Understanding the shared and unique ways artificial and human systems utilize context is therefore a promising direction in enhancing performance in both realms.

## METHODS

All methods were carried out in accordance with relevant guidelines and regulations.

### Stimuli

Stimuli used in this study were photographs of objects against a white background. The objects were selected with a white background to remove confounding background variables and standardize background luminance. Images were obtained from a dataset by Brady et al., (2008), as well as from google image search. Stimuli were 375 × 375 pixels. A total of 730 images were used in this experiment. This was composed of 146 object categories, where each object had five different exemplars (e.g., five different tractors). The 146 object categories were then grouped into pairs of objects that belong to the same context, of which there were 73 contexts. Two contexts, and their corresponding four object categories, were removed from analysis due to their unusually high context and category similarity ratios in VGG CNN analysis - over three standard deviations above the mean. The remaining 71 contextually related pairs of objects were used in all remaining CNN related analyses.

### Human Behavioral Experiments

#### Participants

Participants were recruited online via Prolific (https://www.prolific.co/). Participants self-reported they had normal or corrected to normal vision, fluent in English, and were located in the USA. Participants were financially compensated for their time. All study procedures were approved by the Institutional Review Board of Fordham University. Informed consent was obtained from all participants. In Experiment 1, there were a total of 34 participants (17 females, mean age 36.67, 21-68 range). Two participants were excluded from analysis for not performing the task; in Experiment 2 there were a total of 20 participants (10 females, mean age 34.9, 20-58 range); and in Experiment 3 there were a total of 22 participants (8 females, mean age 28.14, 18-50 age range). Two participants were excluded from analysis in this experiment for not performing the task.

#### Procedure

All experiments were presented using Psychopy software (Peirce et al., 2019) and hosted through the Pavlovia website (https://pavlovia.org). Participants were only permitted to participate in the experiment from a desktop/laptop computer (i.e., no mobile devices).

In all three experiments, a trial began with a fixation cross for 100ms that remained on the screen for the entire trial. Afterwards, two pictures of objects were presented side by side until the participant responded, or up to 1000ms. Participants had up to 3s to make a response. Before the participants began the experiment, they were given 16 practice trials. Participants were asked to respond as quickly and as accurately as they could.

The pair of objects presented were either of the same context or of different contexts. The experiment involved a total of 292 trials, which consisted of four presentations of each object (using four different exemplars). Two trials of each object were presented with a contextually related object, and two trials were presented with a contextually unrelated object (swapped from other contexts). Thus, half of the trials depicted contextually related objects (with two trials per context), and half of which depicted contextually unrelated objects. Specific exemplars of objects were balanced across the conditions across participants.

In Experiment 1, participants were asked to rate how related they found the pair of objects. They responded using a 6-point scale from 1: very dissimilar contexts to 6: very similar contexts. The scale was present on the screen during the duration of the experiment and participants responded using keys 1-6.

In Experiment 2, we wanted to use a paradigm that more accurately reflected reaction time differences across the trials. To accomplish this, participants were asked to make a two alternative forced choice and judge whether the two objects were of the same context (key s) or were of different contexts (key d). Instructions were displayed on the screen for the duration of the experiment. Experiment 2 also included ten catch trials in which two identical objects were presented and the participant had to respond that they were of the same context. This was to increase the quality of data collection.

In Experiment 3, we wanted to use a task more closely related to what a CNN might be picking up on - those objects appear together in the same scene. To accomplish this, in this last study we asked participants to judge whether the two objects would be found in the same photograph (‘s’ for same; ‘d’ for different). Instructions were displayed on the screen for the duration of the experiment. Like Experiment 2, catch trials were also included in this experiment to increase the quality of data collection.

#### Analysis

In all three experiments, we averaged participants’ responses and reaction time for each context (e.g., bike riding) in the related trials (e.g., bike-helmet) and in the unrelated trials (bike-fork). We then ran paired t-test analyses in all experiments to examine whether there were significant differences between related and unrelated responses, both in terms of the relatedness response (depending on the experiment and task) and its reaction time (RT).

### Neural Network Analysis

We focused our study on the VGG16 convolutional neural network, used for its high performance in computer vision and simplicity as an analog to biological neural networks (Simonyan & Zisserman, 2014). We focused on the network as pretrained on the ImageNet data set (Deng et al., 2009; Russakovsky et al., 2015), selected for its wide usage in computer vision due to its large size and its diversity of object classes. Network implementation and analysis was conducted using the Python PyTorch library (Paszke et al. 2019).

The VGG16 network consists of 31 total Layers, each with 64 to 512 units to represent the image; the top layer (layer 31) contains 1000 units, corresponding to the 1000 object categories in ImageNet. Unit responses were extracted at the layers at the end of each processing block of the CNN architecture, where each block begins with a convolution and ends with rectification or max pooling. For VGG16, we studied thirteen layers, specifically: layers 0, 4, 6, 8, 9, 11, 13, 16, 18, 20, 23, 25, and 27. At each layer, representational similarity of image pairs were computed using Pearson’s Correlation of unit responses.

Similarity ratios were computed for each category and context. Specifically for categories, similarities were pooled among image pairs within each category and among image pairs with one image in-category and one image out-of-category. These relative similarities were measured by *SimRatio*^*c*^, computed as follows:

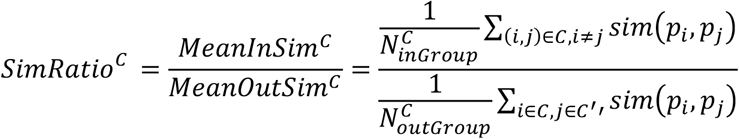

*p*_*i*_,*p*_*j*_ are two images to be compared; in the numerator we consider every pair of distinct images in the category C, i.e., (*i, j*) ∈ *C, i* ≠ *j*; in the denominator, one image is in the category and the other is outside the category, i.e., *i* ∈ *C, j* ∈ *C*′. For several categories C, a set of confounds were removed from the corresponding outside set *C*′, designated for those objects that share a super ordinate category, e.g., a bike helmet was not considered in the “out” group for football helmet. In both the numerator and denominator, the average similarity ratio is computed by dividing the summed ratio by the total number of image pairs in the summation, 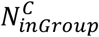 and 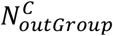. This ratio *SimRatio*^*c*^ was computed for each layer to measure the evolution of category representations through the network layers.

Along the same lines, similarities were pooled among image pairs within each context and among image pairs with one image in-context and one image out-of-context. These relative similarities were measured also by *SimRatio*^*c*^ as computed above. Now, *C* denotes a context rather than a category. Again, a set of confound contexts are removed from *C*’ when computing the denominator. For example, there were multiple object pairs from the kitchen context (e.g., pot – oven mitt; oven – fridge) that would not be considered in the “out” category for one another. For the context ratio, the in-context pairs considered in the numerator exclude all pairs in the same category. For both category and context comparisons, a ratio of 1 indicates no difference in pictures inside and outside the group.

For example, we can consider five easel images representing the “easel” category, five palette images representing the “palette” category, and both categories representing the “art” context. Our category ratio for easel is obtained by dividing the average similarity of distinct easel pictures in-category by the average similarity of easel pictures to non-easel pictures. Our context ratio for art is obtained by dividing the average similarity of easel pictures to palette pictures by the average similarity of easel-or-palette pictures to any other picture in our data set.

Two out of the 73 contexts revealed a context ratio that was several orders of magnitude above the ratios of the other categories. These were removed from further analysis, to assure that these two categories are not driving the results. Thus, all following analyses include 71 contexts.

We repeated our category and context representation analyses on several additional CNN models. We studied additional non-recurrent architectures, VGG 16 and Alexnet, as well as recurrent models Resnet50, Resnet152, and GoogLeNet, all pretrained for object recognition using ImageNet (He et al., 2016; Krizhevsky et al., 2012; Simonyan & Zisserman, 2014; Szegedy et al., 2015). We also studied several architectures pretrained for scene recognition using Places365 (Zhou et al., 2018) For each network, the layer containing the maximum similarity ratio for category and context was studied. (In each network, the same layer had the highest ratio for both context and category.) Recurrent networks were compared to non-recurrent networks and Imagenet-trained networks were compared to Places365-trained networks through T-tests on the average similarity ratio for each of the 142 categories and for each of the 71 contexts.

### Human behavior - CNN correlations

To examine whether the representation of contextual information in CNNs related to human behavior, we computed Pearson correlation between the behavioral results and the CNN results. Specifically, we examined only those trials in which the two objects were related since we investigated the inherent nature of contextual associations. Two behavioral measures were used: the relatedness response and the RT for each of the 71 contexts. These behavioral measures were then correlated with the context similarity ratio in the VGG16 CNN network. First, we correlated behavioral performance with the maximum context ratio found at any layer (typically found at layer 27 for all but one context, which had the maximum ratio at layer 25). We then also examined the relationship between behavior and the representation of each layer of VGG16, and correlated with behavior with the context similarity ratio extracted for each layer of VGG16. The correlations were computed separately for each behavioral experiment.

## Acknowledgments

This work was supported by an interdisciplinary grant from the Office of Research at Fordham University.

## Author Contributions

E.A. designed the project. E.A. & S.B. designed the behavioral studies, collected the behavioral data, and analyzed the data. D.L. & E.R. analyzed the CNN data. E.A., S.B., & D.L. wrote the paper. E.R. contributed to writing the CNN methods. All authors reviewed the manuscript.

The authors declare no competing interests.

## SUPPLEMENTAL MATERAL

**Supplemental Figure:**
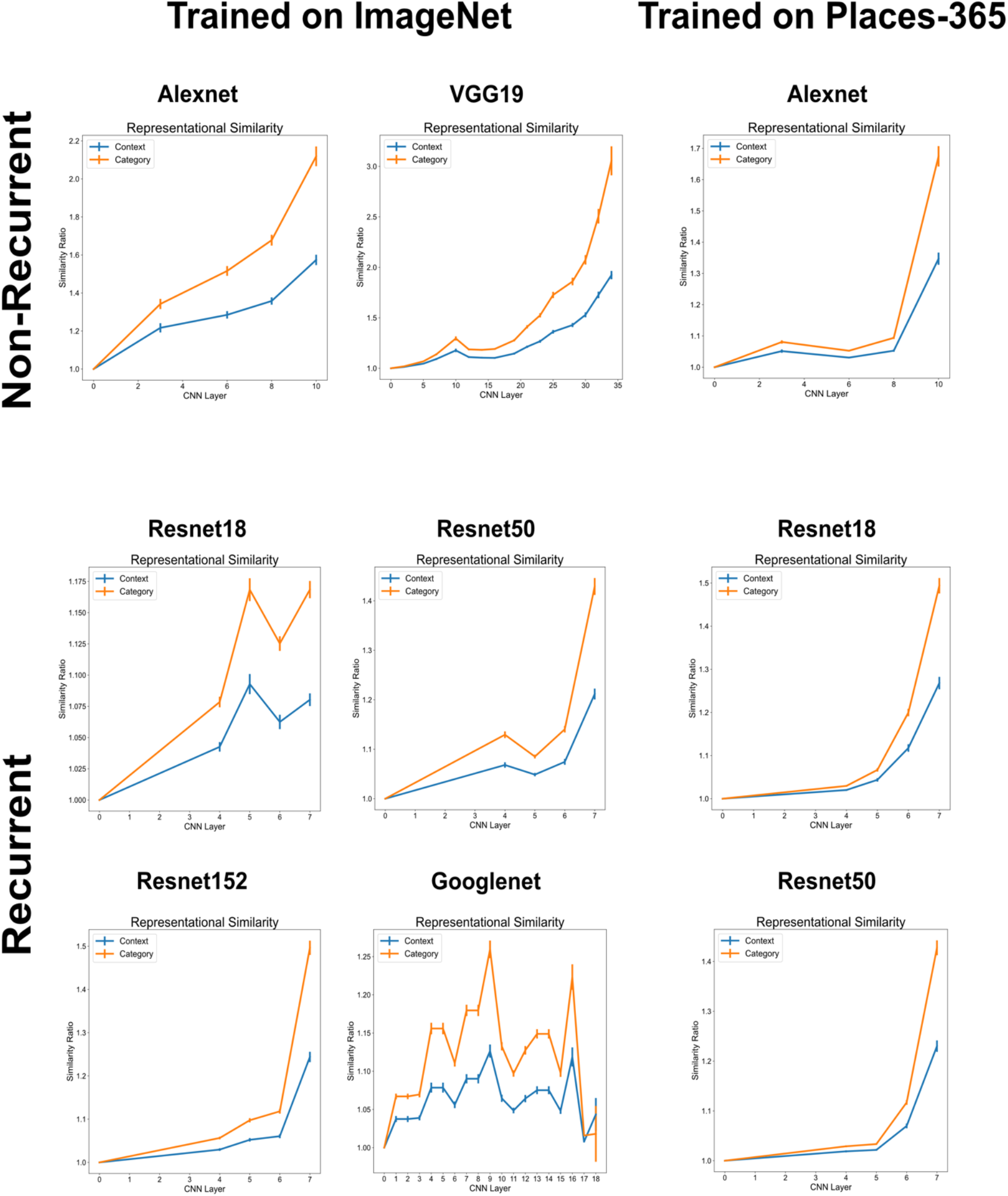
Each graph represents the similarity ratio for category (orange) and context (blue) in each of the networks tested. All networks show a significant effect of contextual associations.

